# Nanoscopic tau aggregates in Parkinson’s disease

**DOI:** 10.1101/2025.08.28.672923

**Authors:** Florence Layburn, Dorothea Boeken, Yu P. Zhang, Kristy Halliday, Dan Rodgers, Lakmini Kahanawita, Bina Patel, Annelies Quaegebeur, Caroline H. Williams-Gray, David Klenerman

## Abstract

Post-mortem tau pathology is frequently observed in Parkinson’s disease (PD) using immunohistochemistry (IHC) to measure large inclusions, however, small protein aggregates that precede inclusions are considered a major driver of toxicity in neurodegenerative disease. We aimed to uncover the distribution of nanoscopic aggregates across six brain regions in post-mortem tissue from 14 PD and 15 controls using the single-molecule pull-down assay (SiMPull). In the hippocampus and amygdala, tau IHC and SiMPull were associated with advanced age in controls and dementia status in PD. Despite negligible tau IHC-labelled aggregates in the putamen, we identified a unique population of high-intensity nanoscopic tau aggregates for a subset of PD cases using SiMPull, ranging from 10–1,000 epitopes per aggregate and 30–1,000 nm in length. Previous evidence linking nigrostriatal tau pathology and motor deficits indicates that the nanoscopic tau aggregates identified in this study may contribute to striatal dysfunction in PD.

## Main

Parkinson’s disease (PD) manifests as a movement disorder during life, however, post-mortem neuropathology is defined by two key features. First is the loss of dopaminergic neurons from the substantia nigra in the midbrain, a component of the basal ganglia movement-control circuitry (1, 2). Cell death in the substantia nigra directly translates to the cardinal motor symptoms of PD, such as slowness of movement and rigidity (3). The second criterion for PD diagnosis is the presence of Lewy Bodies; large, insoluble protein aggregates primarily composed of alpha-synuclein (αSyn) and distributed throughout the brain according to the Lewy Body pathology Braak stages (4–6). Post-mortem neuropathology has informed the development of biomarkers to help support the diagnosis of PD during life, for example DaTScans to assess the loss of dopaminergic nerve terminals in the striatum, and the explosion of αSyn-based fluid biomarkers currently in preclinical development (7–11). Our understanding of PD neuropathology is overwhelmingly αSyn-centric, however, this is not the only protein to aggregate in the PD brain.

For many years, post-mortem tau pathology has also been observed in PD. Tau is a microtubule-associated protein that becomes hyperphosphorylated and forms inclusions such as neurofibrillary tangles in the brains of people with Alzheimer’s disease and other tauopathies (12, 13). There are surprisingly few studies on the significance of tau in PD considering that approximately 50% of PD-dementia cases display post-mortem tau pathology (14); there is an association between genetic variation in MAPT, the gene encoding tau, and increased risk of parkinsonism (15); and there are several parkinsonian disorders affecting the nigrostriatal pathway, such as progressive supranuclear palsy, where the primary pathology is tau (16, 17). Furthermore, a recently published study identified an association between nigrostriatal tau pathology and motor deficits in a cohort of PD and non-PD individuals, suggesting that tau may contribute to dopaminergic degeneration (18). Despite the overlap of pathology in neurodegenerative diseases, we still do not know if tau aggregation is merely co-incident and independent of PD disease progression, or co-pathological and actively contributing to cell death in PD.

Currently, neuropathological analysis depends on detection of large insoluble inclusions which are visible by immunohistochemistry (IHC). However, Lewy Bodies are poorly associated with neurological dysfunction and mounting evidence suggests that small, intermediate protein aggregates— which are undetectable using IHC due to their nanoscopic size range— are the true drivers of disease for a range of neurological disorders (19–21). For example, small aggregates are more potent at inducing neurotoxicity, and more efficient at spreading between cells to seed further aggregation when compared to fibrils and inclusion bodies (22, 23). To overcome the barriers to studying nanoscopic aggregates, our group uses the single-molecule pull-down assay (SiMPull) and fluorescence imaging to characterise protein aggregates from human samples in terms of size, concentration, and composition without the need for excessive purification or denaturing conditions that would disrupt endogenous aggregate structures. We have used SiMPull to measure protein aggregates from blood, cerebrospinal fluid, and brain samples; for example, identifying diffusible αSyn aggregates in PD brain tissue capable of inducing inflammation and membrane permeabilisation (20, 24, 25).

In this study, we aimed to improve our understanding of tau aggregate distributions for the full range of aggregate sizes in a cohort of 14 PD and 15 controls (CON). To capture the breadth of pathology in the PD brain, we studied six brain regions; from earliest to latest affected according to the Lewy Body Braak stages, including the substantia nigra (SN), amygdala (AMYG), hippocampus (HIP), putamen (PUT), frontal cortex (FC) and occipital cortex (OC). We measured tau aggregates extracted from fractionated post-mortem brain tissue— henceforth referred to as nanoscopic aggregates— using SiMPull, and measured tau inclusions in formalin-fixed, paraffin-embedded tissue sections using IHC. We identified a disease-specific trend of higher intensity nanoscopic tau aggregates in the basal ganglia using SiMPull, implicating tau in striatal dysfunction in PD.

## Results

### Single-molecule imaging can be used to measure protein aggregates’ abundance, intensity, and size

We used immunohistochemistry (IHC) and single-molecule fluorescence imaging to quantify protein aggregates from post-mortem brain tissue from 14 PD and 15 controls across six brain regions (Figure 1a). The groups were similar in age (PD = 80.9 ± 6.5 years; control = 74.7 ± 12.7 years; two-sided T-test *p*-value = 0.13) and post-mortem interval (PD = 53.2 ± 32.1 hours; control = 53.8 ± 27.1 hours; two-sided T-test *p*-value = 0.96), and there were slightly more females in the control group (6F/9M) compared to PD (3F/11M) (Methods Table 1). Formalin-fixed, paraffin-embedded (FFPE) brain tissue sections were stained using 3, 3’-diaminobenzidine (DAB) IHC to quantify large insoluble aggregates, and fractionated fresh-frozen brain tissue was used for single-molecule pull-down (SiMPull) to measure nanoscopic aggregates (Figure 1b–c). SiMPull is an aggregate-specific assay, due to the use of identical capture and detection antibodies; if a monomer is captured on the surface, it does not have a second epitope to accommodate the detection antibody and therefore does not generate a fluorescent signal (Figure 1c).

**Figure 1.**
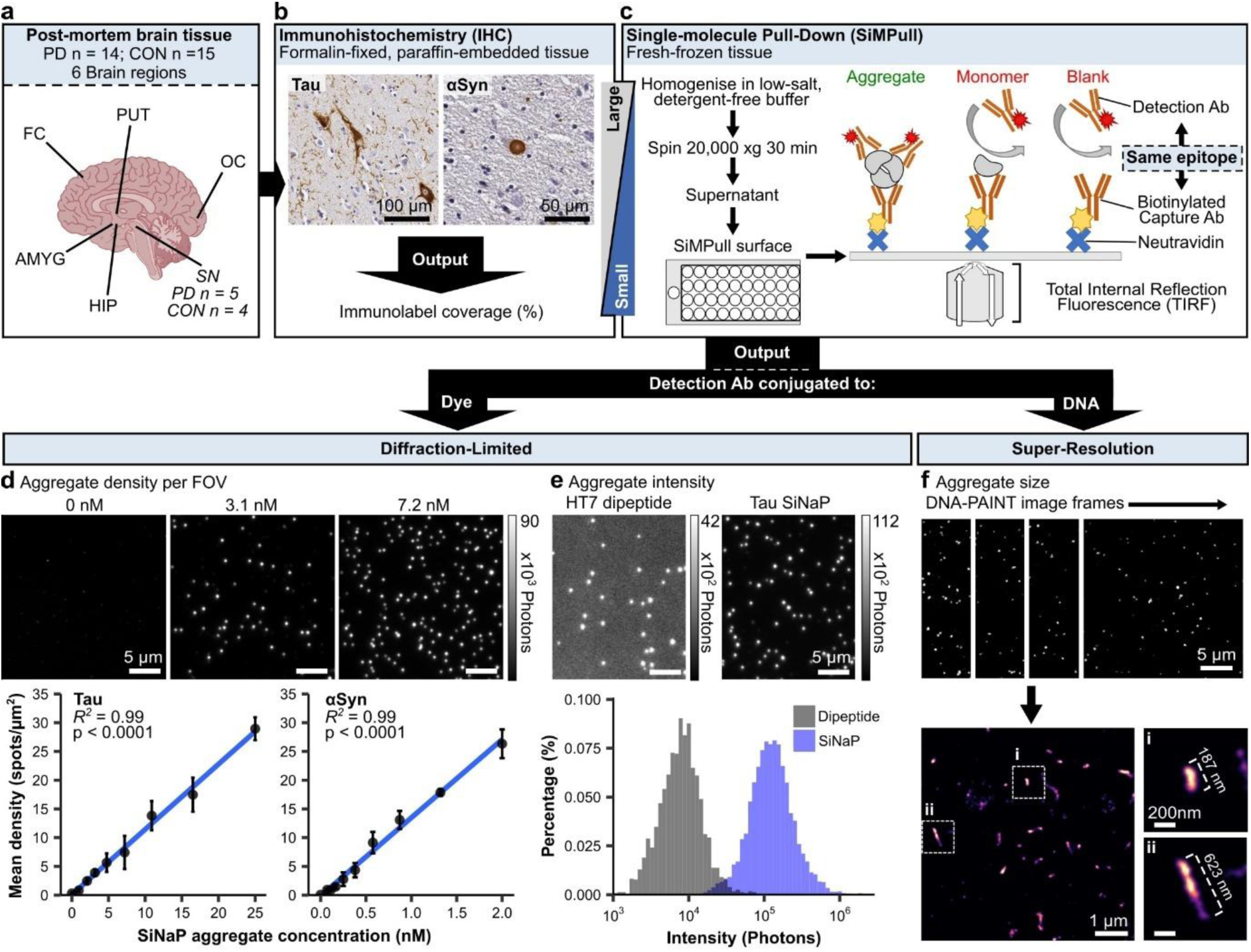
SiMPull workflow and output for characterising protein aggregates. **(a)** Post-mortem brain tissue cohort overview. The SN was highlighted due to the reduced sample size for frozen tissue. **(b)** IHC was used to study large, insoluble inclusion bodies. **(c)** Schematic representation of SiMPull used to detect nanoscopic aggregates. SiMPull excludes monomers from detection by using identical capture and detection antibodies. **(d)** Examples of diffraction-limited imaging using tau SiNaP aggregate standards. Aggregate density was significantly correlated with aggregate concentration for tau and αSyn SiNaP aggregates (Pearson correlation). Each point represents the mean of 12 FOVs and three well replicates, error bars represent mean ± standard deviation. **(e)** “Auto” contrast-adjusted example images of HT7 dipeptide, tau SiNaPs, and corresponding aggregate intensity distributions. **(f)** Example DNA-PAINT images from one FOV are shown; first the individual image frames, then the super-resolved image. Insets **(i)** and **(ii)** feature close-ups of super-resolved phosphorylated tau aggregates from a PD human brain sample. CON = Control; HIP = Hippocampus; AMYG = Amygdala; PUT = Putamen; SN = Substantia nigra; FC = Frontal cortex; OC = Occipital cortex. SiNaP = silica nanoparticle; FOV = Field of view.

**Table 1.** Clinical information for all PD and Control (CON) brain tissue samples. N/A = Not available; AOD = Age of Death; PMI = Post-mortem interval; FTD = Frontotemporal dementia; MND = Motor neuron disease; GI = Gastrointestinal.

| Case | Sex | AOD (years) | PMI (hours) | PD Duration (years) | Lewy Body Braak Stage | Tau Braak & Braak stage | Dementia status | Cause of Death |
| --- | --- | --- | --- | --- | --- | --- | --- | --- |
| PD1 | M | 71 | 64 | 4.4 | VI | II | N | N/A |
| PD2 | M | 85 | 8 | 18.1 | VI | III | Y | PD Dementia |
| PD3 | M | 73 | 57 | N/A | V | II | N | PD |
| PD4 | F | 78 | 68 | 5.2 | V | II | N | Pneumonia |
| PD5 | M | 76 | 60 | 9.7 | V | II | Y | Aspiration Pneumonia |
| PD6 | M | 77 | 101 | 8.5 | V | III | N | Pneumonia |
| PD7 | F | 86 | 52 | N/A | VI | II | N | PD |
| PD8 | M | 86 | 32 | 4.8 | V | III | N | Myocardial Infarction |
| PD9 | M | 84 | 21 | 6.3 | IV | II | N | Pneumonia |
| PD10 | M | 89 | N/A | 11.9 | V | I | Y | Hypertensive heart disease |
| PD11 | M | 88 | 44 | 22.4 | VI | III | N | Pneumonia |
| PD12 | M | 73 | 124 | N/A | VI | II | N | Massive GI bleed |
| PD13 | F | 90 | 52 | N/A | V | III | Y | Vascular dementia |
| PD14 | M | 75 | 8 | 11.5 | VI | III | Y | PD Dementia |
| CON1 | M | 72 | 39 | - | 0 | II | N | Malignant tumor of spinal cord |
| CON2 | F | 65 | 5 | - | 0 | II | N | Idiopathic Pulmonary Fibrosis |
| CON3 | M | 39 | 59 | - | 0 | 0 | N | Metastatic cancer of the colon |
| CON4 | M | 73 | 51 | - | 0 | II | N | Myocardial infarction |
| CON5 | M | 62 | 63 | - | 0 | I | N | Hepatocellular carcinoma |
| CON6 | M | 74 | 74 | - | 0 | II | N | N/A |
| CON7 | M | 88 | 18 | - | 0 | III | N | Acute haemopericardium |
| CON8 | F | 88 | 28 | - | 0 | III | N | Multiorgan failure |
| CON9 | F | 77 | 59 | - | 0 | III | N | Burkitts Lymphoma |
| CON10 | F | 75 | 92 | - | 0 | II | N | Metastatic ovarian Cancer |
| CON11 | M | 87 | 49 | - | 0 | III | Y | Dementia |
| CON12 | F | 86 | 67 | - | 0 | III | N | Idiopathic Pulmonary Fibrosis |
| CON13 | F | 70 | 32 | - | 0 | I | N | N/A |
| CON14 | M | 90 | 56 | - | 0 | II | Y | FTD/MND |
| CON15 | M | 74 | 115 | - | 0 | II | Y | Dementia |

The output of SiMPull depends on the imaging probe; a dye-conjugated detection antibody was used for diffraction-limited imaging, and a DNA-conjugated antibody together with a dye-labelled imaging DNA strand was used for DNA points accumulation for imaging in nanoscale topography (DNA-PAINT) super-resolution imaging. We used diffraction-limited imaging to quantify the density of aggregates per field of view as a measure of abundance (Figure 1d) and aggregate intensity as a measure of size (Figure 1e). To demonstrate the utility of diffraction-limited SiMPull, we used silica nanoparticles (SiNaPs) as aggregate standards, either conjugated with the tau antibody HT7 binding sequence or αSyn monomers. There was a strong linear relationship between SiNaP concentration and SiMPull density for a dilution series of tau and αSyn SiNaPs respectively (Figure 1d). The tau SiNaP intensity distribution was higher than the HT7 antibody binding sequence dipeptide, which can only bind a single detection antibody, demonstrating that larger aggregates with higher numbers of detection antibody binding lead to higher intensity values (Figure 1e).

We used DNA-PAINT to expand on the size information from diffraction-limited SiMPull and measure aggregate lengths on the nanometre scale. DNA-PAINT overcomes the diffraction-limit of light by separating the fluorescent signal over time across thousands of frames, which were subsequently reconstructed to generate the super-resolved image (Figure 1f).

### PD cases display typical neuropathology in FFPE sections, but αSyn SiMPull does not detect PD-specific nanoscopic aggregates

All PD cases in this study met the international criteria for post-mortem diagnosis of PD, displaying neurodegeneration in the substantia nigra (Figure 2a–b) and Lewy Body pathology (Figure 2c–e). As expected, αSyn immunolabel coverage was significantly higher in PD compared to controls in all brain regions except the occipital cortex, which had low staining in both groups (Figure 2c). Typical Lewy Body staining and elongated, neuritic αSyn staining was observed by IHC across the PD samples (Figure 2d–e).

**Figure 2.**
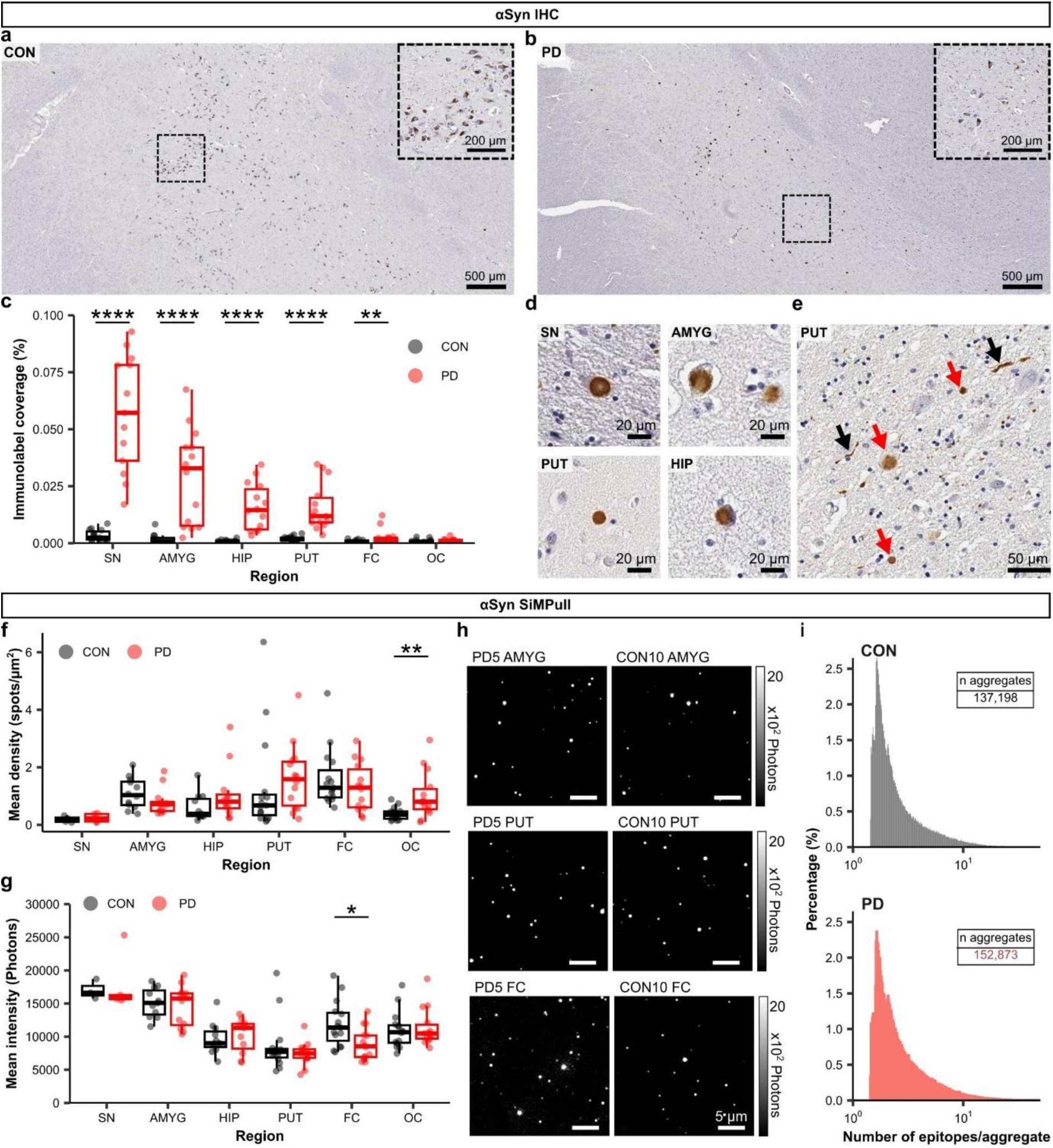
PD cases displayed typical neuropathology in FFPE brain sections, but αSyn SiMPull does not detect PD-specific nanoscopic aggregates. (a–b) Representative images of neuronal loss in the SN in PD compared to CON. **(c)** αSyn IHC coverage was significantly higher in PD (n =14) compared to CON (n = 15) for all brain regions except the OC, which had low staining in both groups (Two-sided Mann-Whitney test, SN *p* < 0.0001; AMYG *p* < 0.0001; PUT *p* < 0.0001; HIP *p* < 0.0001; FC *p* = 0.0023; OC *p* = 0.1583). Each point represents an individual case. **(d–e)** Representative images of αSyn staining in PD brain tissue. **(e)** Depicts different staining morphologies within close proximity in a PUT section, with red arrows highlighting Lewy Bodies as round inclusion-like staining and black arrows highlighting Lewy Neurites as elongated, neuritic staining. **(f)** αSyn SiMPull mean aggregate densities were significantly higher in the PD OC compared to CON (Two-sided Mann-Whitney test, SN *p* = 0.9048; AMYG *p* = 0.1658; PUT *p* = 0.1583; HIP *p* = 0.2115; FC *p* = 0.4253; OC *p* = 0.0079). **(g)** αSyn SiMPull mean aggregate intensities were significantly higher in the CON FC compared to PD (Two-sided Mann-Whitney test, SN *p* = 0.4127; AMYG *p* = 0.4776; PUT *p* = 0.7148; HIP *p* = 0.3164; FC *p* = 0.01375; OC *p* = 0.9145). **(f–g)** Each point represents the mean of 9–12 FOVs and 2–3 independent repeats per case. **(h)** Example αSyn SiMPull images from one PD and one CON case in the AMYG, PUT, and FC. All images were contrast-adjusted to the same range. **(i)** Distribution of epitopes per aggregate for CON and PD compiled for all brain regions, plotted on a log scale. CON = Control; SN = Substantia nigra; AMYG = Amygdala; HIP = Hippocampus; PUT = Putamen; FC = Frontal cortex; OC = Occipital cortex; FOV = Field of view.

We used the αSyn C-terminus binding antibody Syn211 that detects most forms of αSyn for SiMPull to measure nanoscopic αSyn aggregates and compare with Lewy Body deposition (26). SiMPull aggregate densities were within the lower range of detection as compared to αSyn SiNaPs in Figure 1b, mostly below 2 spots/µm^2^ for all brain regions, and there was a significant increase in the PD occipital cortex compared to controls (Figure 2f). Aggregate intensities were also low, although there was a significant increase in mean aggregate intensity in the control frontal cortex compared to PD (Figure 2g). In light of the low signal from SiMPull signal, we confirmed that Syn211 binds αSyn in the fractionated brain samples (Supplementary Figure 1a), and obtained similar SiMPull results with five alternative αSyn antibodies (Supplementary Figure 1b–d).

We probed the intensity values further by calculating the approximate number of epitopes per aggregate. To do so, we used another total αSyn antibody SOY1 to capture αSyn monomers on the surface and detected using Syn211 (Methods Figure 1). This analysis revealed that 88.8% of aggregates for controls and 87.8% of aggregates for PD were composed of fewer than five epitopes, and there were no clear differences between the size distributions of PD and controls (Figure 2i). Due to the low abundance and small size of objects detected by αSyn SiMPull, we did not consider these species disease-specific, and continued to investigate tau protein aggregates in the PD brain.

### Tau SiMPull reveals two sub-populations of nanoscopic aggregates, and higher intensity aggregates in the PD putamen

To investigate the distribution of tau pathology in PD, we first used IHC to quantify tau immunolabel coverage (Figure 3a). There was a wide range of tau staining levels in the hippocampus and amygdala across the whole tissue cohort, although no significant differences between PD and controls (Figure 3a). For future analysis, control cases were subclassified into Path-Positive and Path-Negative based on tau staining levels in the hippocampus (Figure 3a). There was a significant increase in tau staining in the PD substantia nigra, although absolute coverage levels were low compared to the hippocampus and amygdala, which had visibly higher staining coverage compared to the other regions (Figure 3b).

**Figure 3.**
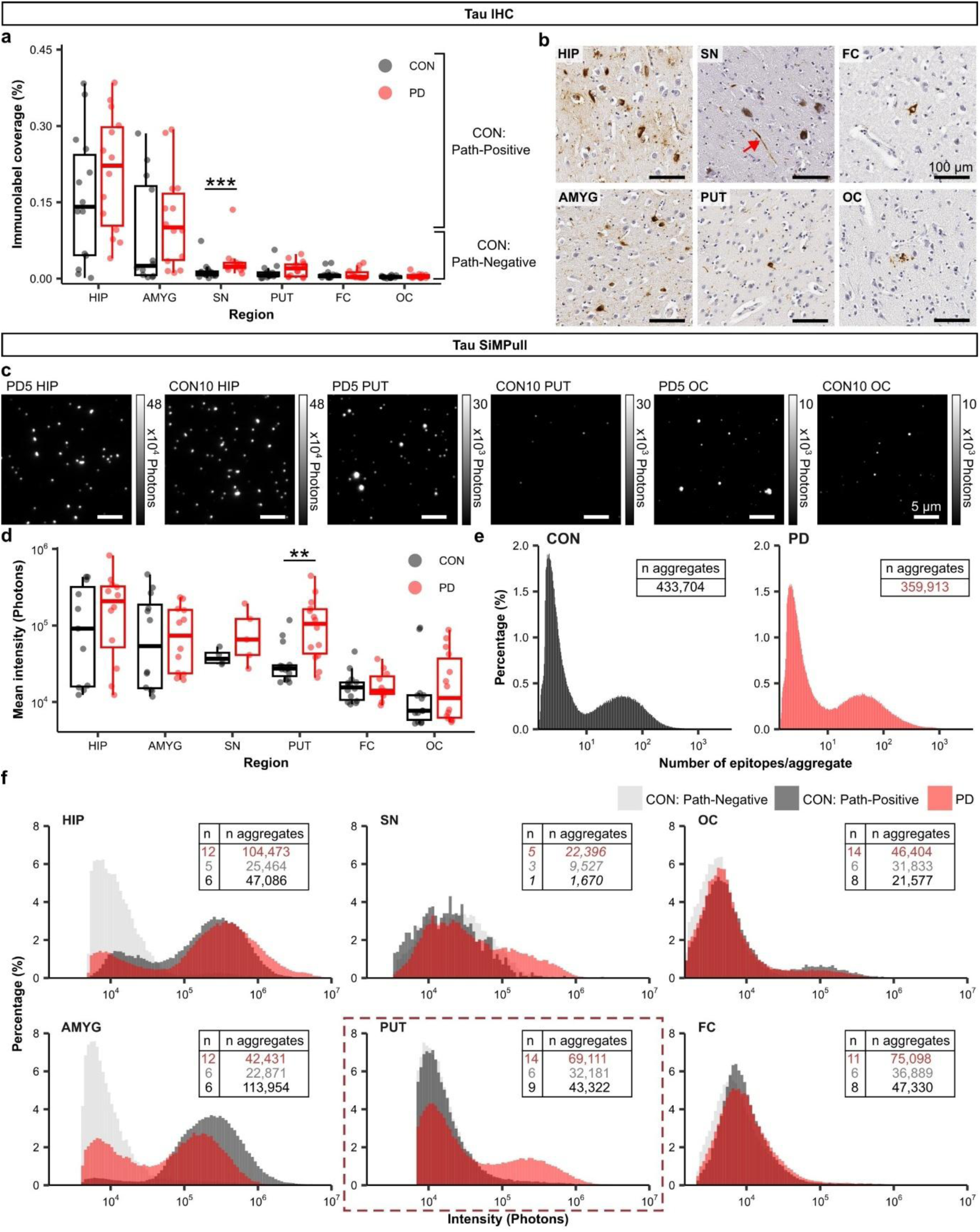
Tau SiMPull intensities are higher in the PD putamen compared to CON. **(a)** Tau IHC coverage in PD (n = 14) compared to CON (n = 15) shows a significant increase in PD in the SN (Two-sided Mann-Whitney test, HIP *p* = 0.4773; AMYG *p* = 0.2116; SN *p* = 0.00044; PUT *p* = 0.1456; FC *p* = 0.5045; OC *p* = 0.3314). Each point represents an individual case. **(b)** Representative images of tau staining. The red arrow highlights tau staining in the SN as opposed to dark pigmented cells also visible. **(c)** Example tau SiMPull images from one PD and one CON case across the HIP, PUT, and OC. Each pair of images was contrast-adjusted to the same range. **(d)** Tau SiMPull mean aggregate intensity, plotted on a log scale. Aggregate intensities were significantly higher in the PD PUT compared to CON (Two-sided Mann-Whitney test, HIP *p* = 0.6075; AMYG *p* = 0.8428; SN *p* = 0.4127; PUT *p* = 0.0013; FC *p* = 0.9358; OC *p* = 0.3761). Each point represents the mean of 9–12 FOVs and 2–3 independent repeats per case. **(e)** Distribution of epitopes per aggregate for CON and PD respectively compiled for all brain regions, plotted on a log scale, revealing two populations of aggregate sizes. **(f)** Tau aggregate intensity distributions for Path-Negative CON, Path-Positive CON, and PD across all brain regions. The PUT is highlighted to demonstrate the higher intensity distribution in PD. For each region, “n” refers to the number of cases, and “n aggregates” refers to the number of aggregates detected by SiMPull per group. SN sample numbers are italicised to highlight the lower n compared to other brain regions. CON = Control; HIP = Hippocampus; AMYG = Amygdala; PUT = Putamen; SN = Substantia nigra; FC = Frontal cortex; OC = Occipital cortex; FOV = Field of view.

We followed up IHC analysis with tau SiMPull using the total-tau antibody HT7 and observed similar trends in the hippocampus and amygdala, with high intensity aggregates for both PD and controls in these regions (Figure 3c–d). Interestingly, there was a significant increase in aggregate intensities in PD compared to controls in the putamen that was not previously observed using IHC (Figure 3d). There were no significant differences in aggregate density between controls and PD in any of the brain regions (Supplementary Figure 2a).

We used the HT7 dipeptide to calculate the approximate number of epitopes per aggregate and observed two distributions in both PD and controls, reflecting two aggregates populations of 1–10 and 10–1000 epitopes respectively (Figure 3e). This prompted us to inspect the underlying intensity distributions for PD, Path-Positive controls and Path-Negative controls per brain region (Figure 3f). The intensity distributions revealed there were both low and high-intensity aggregates in PD and Path-Positive controls in the hippocampus and amygdala; there were predominantly low-intensity aggregates in the frontal and occipital cortices for all groups; and there were predominantly low-intensity aggregates in Path-Negative controls in all brain regions. In the putamen, the high-intensity aggregate population was only present in PD.

**Methods Figure 1.**
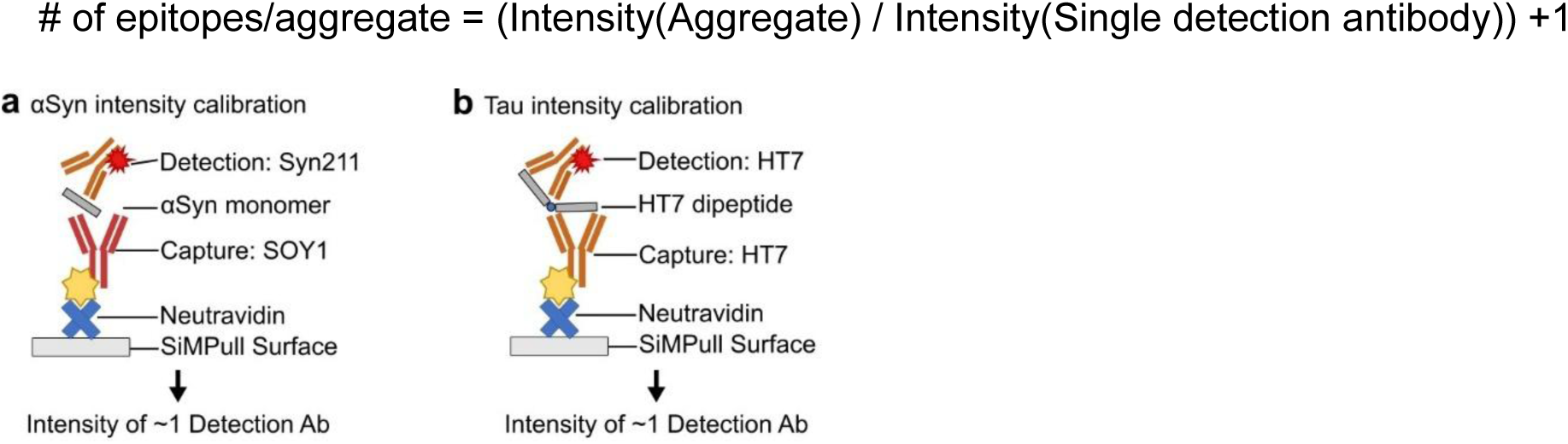
Intensity calibrants to calculate the approximate number of epitopes per aggregate. Schematic representation of **(a)** αSyn and **(b)** tau intensity calibrants used to measure the intensity of a single detection antibody for calculating the approximate number of epitopes per aggregate.

### Phosphorylated tau aggregates are higher intensity in PD putamen compared to CON

We repeated SiMPull using the phosphorylated tau (pTau) antibody AT8, which is commonly used for neuropathological diagnosis of tauopathies. Again, high-intensity aggregates were observed in the hippocampus and amygdala for both PD and controls (Figure 4a–b). Furthermore, mean aggregate intensities were correlated with tau SiMPull and pTau SiMPull (Supplementary Figure 2b). Mean aggregate intensities were significantly higher in PD in the putamen and substantia nigra, although the latter had a reduced sample size and did not reach statistical significance, and there was a modest increase in intensity in the PD occipital cortex (Figure 4b). Although aggregate levels were lower in the occipital cortex compared to other brain regions, we also observed a significant increase in aggregate density in PD in this region (Figure 4c). Unlike tau SiMPull, pTau aggregate intensities displayed a single distribution for each group (Figure 4d). There were overlapping distributions for PD and Path-Positive controls in the hippocampus and amygdala, and a distinct shift towards higher pTau aggregate intensities for PD in the putamen and substantia nigra (Figure 4d). In the occipital cortex, the difference in intensities observed in Figure 4c was explained by the predominantly low-intensity aggregates in Path-Negative controls, however pTau aggregate intensities were similar between Path-Positive controls and PD (Figure 4d).

**Figure 4.**
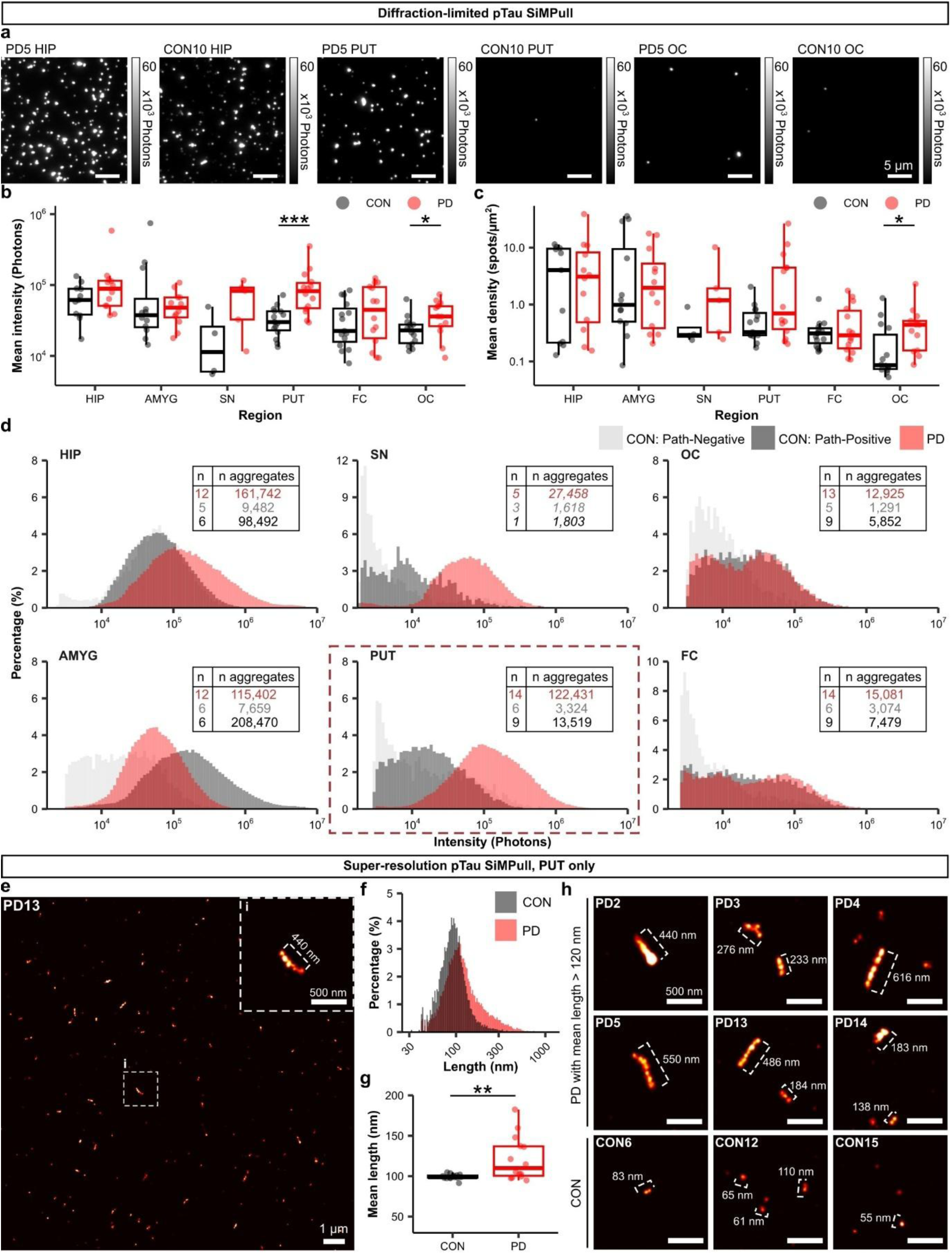
pTau aggregates are higher intensity in PD PUT compared to CON, corresponding to lengths of 30–1,000 nm. **(a)** Example pTau SiMPull images from one PD and one CON case across the HIP, PUT, and OC. All images were contrast-adjusted to the same range. **(b)** pTau SiMPull mean aggregate intensity for PD (n = 14) and CON (n = 15), plotted on a log scale. Aggregate intensities were significantly higher in the PD PUT and OC (Two-sided Mann-Whitney test, HIP *p* = 0.2115; AMYG *p* = 0.5899; SN *p* = 0.1111; PUT *p* = 0.00022; FC *p* = 0.2703; OC *p* = 0.04817). **(c)** pTau SiMPull mean aggregate density, plotted on a log scale. Aggregate densities were significantly higher in the PD OC compared to CON (Two-sided Mann-Whitney test, HIP *p* = 0.9759; AMYG *p* = 1; SN *p* = 0.1905; PUT *p* = 0.1121; FC *p* = 0.9486; OC *p* = 0.01448). **(b–c)** Each point represents the mean of 9–12 FOVs and 2–3 independent repeats per case. **(d)** Tau aggregate intensity distributions for Path-Negative CON, Path-Positive CON, and PD across all brain regions. The PUT is highlighted to demonstrate the significant shift towards higher intensity aggregates in PD. For each region, “n” refers to the number of cases, and “n aggregates” refers to the number of aggregates detected by SiMPull per group. SN sample numbers are italicised to highlight the lower n compared to other brain regions. **(e)** Example super-resolution image of PD13 from a subset of the FOV. **(f)** Length distributions of super-resolved pTau aggregates in the PUT for PD and CON, ranging from 30–1,000 nm. **(g)** Aggregate lengths were significantly higher in PD compared to CON (Two-sided Mann-Whitney test, *p* = 0.0099). Each point represents the mean of 4 FOVs per case. **(h)** Example images of long aggregates from the highest six PD cases and representative images of short aggregates from CON cases. A subset of the FOV was used for demonstration purposes and the fire colour preset was used to represent the density of localisations. CON = Control; FOV = Field of view; HIP = Hippocampus; AMYG = Amygdala; PUT = Putamen; SN = Substantia nigra; FC = Frontal cortex; OC = Occipital cortex;

We identified pTau aggregates in the putamen ranging from 30–1,000 nm using super-resolution imaging (Figure 4e–f). There was a significant increase in aggregate length in PD, driven by a subset of PD cases with longer aggregates (Figure 4g). Examples of long pTau aggregates in the six highest PD cases are demonstrated in Figure 4h, accompanied by representative images of smaller aggregates that were present in both PD and controls.

### SiMPull tau aggregate intensities are correlated with IHC tau coverage the hippocampus and amygdala, but not PD putamen

Next, we investigated the relationship between tau IHC coverage and tau SiMPull aggregate intensities. We started by classifying the nanoscopic aggregates from SiMPull as “dim” or “bright” and calculated the percentage of bright aggregates per case, revealing that a combination the two aggregate populations could be detected within an individual (Figure 5a– b). The brightness trends were consistent with tau SiMPull mean intensities and there was a significant increase in PD putamen compared to controls (Figure 5b).

**Figure 5.**
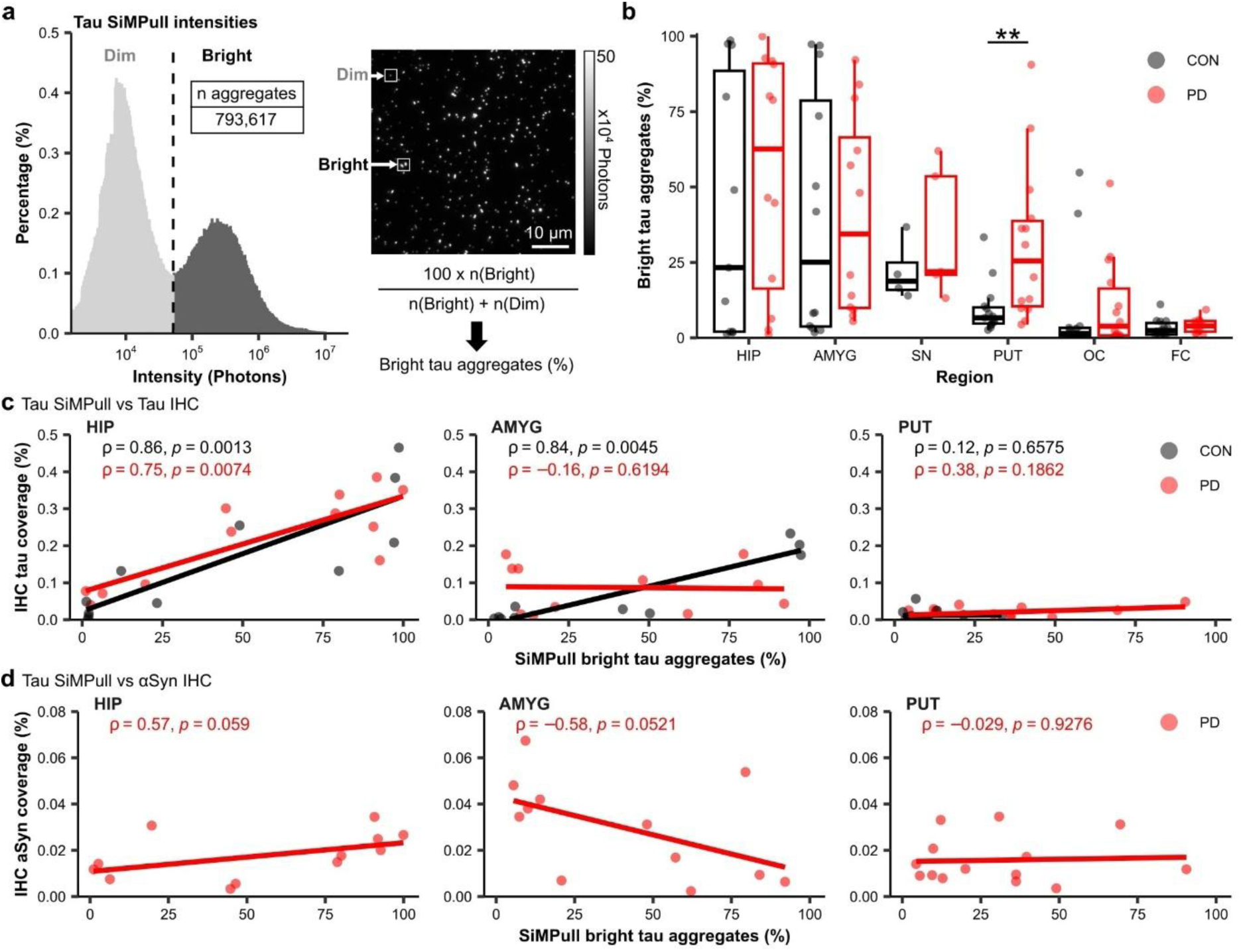
Tau SiMPull is correlated with tau IHC in CON, but not in PD PUT. **(a)** Pooled tau SiMPull intensity distribution for all regions in PD (n = 14) and CON (n = 15). Aggregates were classified as “dim” or bright”, and the percentage of bright aggregates for each case was calculated. A representative image of dim and bright aggregates from tau SiMPull is shown. **(b)** There was a significant increase in bright tau aggregates in PD in the PUT (Two-sided Mann-Whitney test, HIP *p* = 0.6075; AMYG *p* = 0.5899; SN *p* = 0.5556; PUT *p* = 0.0023; FC *p* = 0.3173; OC *p* = 0.4544). Each point represents the mean of 9–12 FOVs and 2–3 independent repeats per case. **(c)** Tau SiMPull bright aggregates were positively correlated with IHC tau coverage in CON HIP and AMYG, but not PUT (Spearman correlation). Tau SiMPull bright aggregates were positively correlated with IHC tau coverage in PD HIP, but not AMYG or PUT (Spearman correlation). **(d)** Tau SiMPull bright aggregates and αSyn IHC coverage were correlated in PD HIP and AMYG, although these did not reach significance (Spearman correlation). **(c–d)** Each point represents the mean per case. CON = Control; HIP = Hippocampus; AMYG = Amygdala; PUT = Putamen; SN = Substantia nigra; FC = Frontal cortex; OC = Occipital cortex; FOV = Field of view.

There was a significant correlation between tau SiMPull and tau IHC in the hippocampus for both controls and PD (Figure 5c). There was also a significant correlation between tau SiMPull and tau IHC in the amygdala for controls. However, there was no significant correlation between tau SiMPull and tau IHC for PD in the amygdala, or for PD and controls in the putamen. The latter can be explained by the low signal for controls in the putamen, however, this analysis highlighted the presence of bright nanoscopic tau aggregates in PD that were undetectable by IHC.

In PD, there was a positive correlation between tau SiMPull and αSyn IHC coverage in the hippocampus, and a negative correlation in the amygdala, although neither relationship reached statistical significance (Figure 5d). Due to low signal, there was no significant correlation between SiMPull and IHC in substantia nigra, frontal cortex, and occipital cortex brain regions (Supplementary Figure 3).

### Tau pathology is associated with age in controls and dementia status in PD

We used principal component analysis (PCA) and k-means cluster analysis to summarise the different pathology trends within this cohort (Figure 6a). The PCA input variables included αSyn IHC coverage (%), tau IHC coverage (%), tau SiMPull bright aggregates (%) and pTau SiMPull mean intensity (photons) from the hippocampus, amygdala, and putamen. αSyn IHC coverage and tau IHC from the substantia nigra were also included, however SiMPull results from the substantia nigra were excluded due to low sample size. Results from the frontal and occipital cortices were excluded due to low signal across all markers. We identified four clusters within the cohort, comprised of two control and two PD sub-groups (Figure 6a). We plotted the pathology markers for each PCA group to reveal: Group 1 controls had low pathology in all brain regions; Group 2 controls had high tau pathology predominantly in the medial temporal lobe (including the hippocampus and amygdala); Group 3 PD cases had αSyn pathology and moderate tau pathology in the medial temporal lobe; Group 4 PD cases had αSyn pathology, as well as high tau pathology in the medial temporal lobe and putamen (Figure 6b). The pathology trends for the individual cases in each group are represented in Supplementary Figure 4.

**Figure 6.**
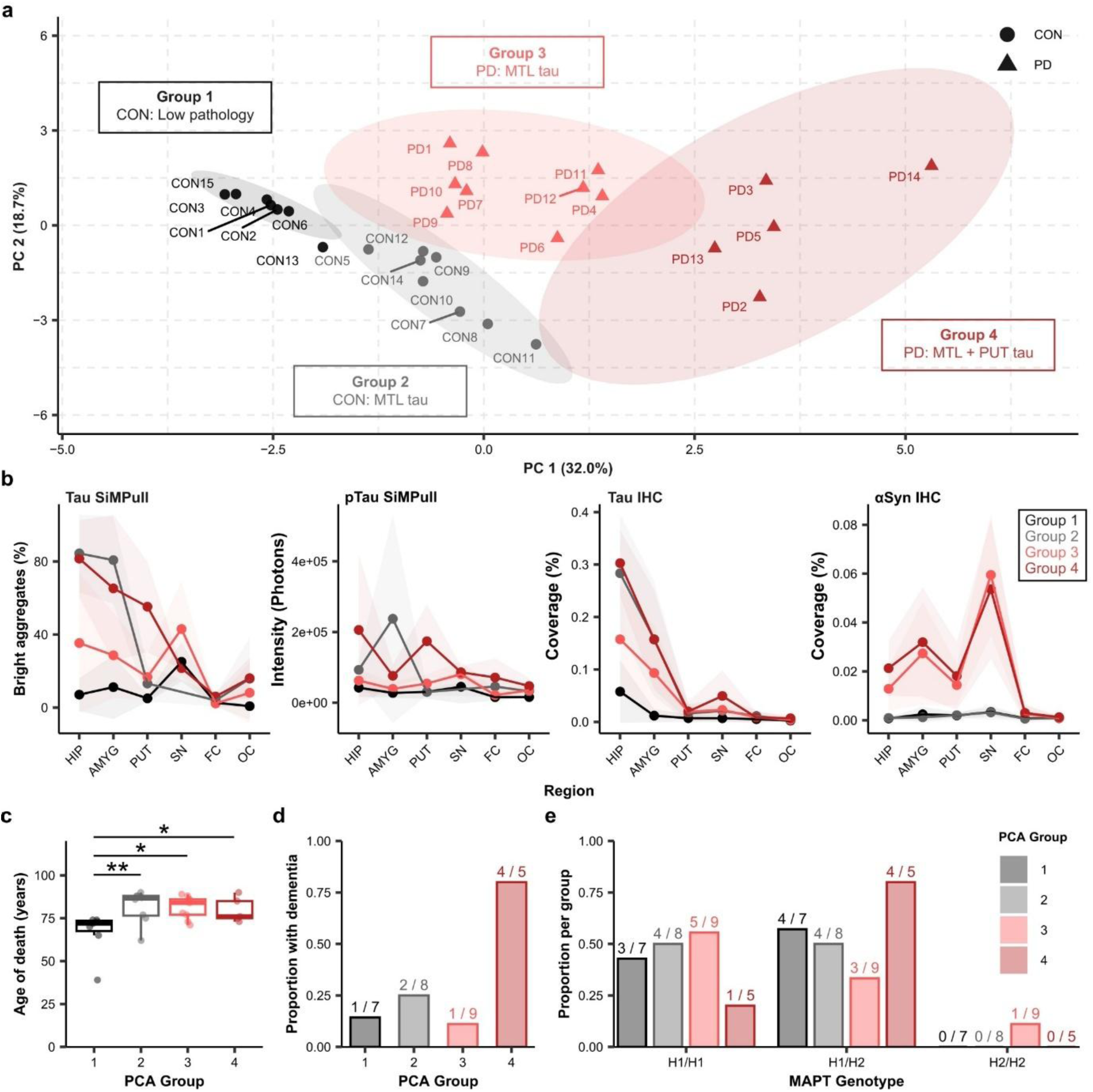
Tau pathology is associated with age in CON and dementia status in PD. **(a)** Principal component analysis (PCA) shows the separation of the cohort into four clusters along PC1 (32.0% variance) and PC2 (18.7% variance). Ellipses indicate confidence interval = 0.95 of groups. MTL = Medial temporal lobe, referring to the HIP and AMYG. **(b)** Summary of pathology for each PCA group, each point represents the group mean and the shaded area represents the standard deviation. **(c)** Age of death was significantly lower in Group 1 compared to the other groups (Dunn’s multiple comparison test, Group 1 vs Group 2 *p* = 0.0099; Group 1 vs Group 3 *p* = 0.0120; Group 1 vs Group 4 *p* = 0.0393). **(d)** Group 4 had the highest proportion of cases with a dementia diagnosis. **(e)** There was an even distribution of H1/H1 and H1/H2 MAPT genotype for Groups 1, 2 and 3. The majority of cases in Group 4 had an H1/H2 MAPT genotype. CON = Control; HIP = Hippocampus; AMYG = Amygdala; PUT = Putamen; SN = Substantia nigra; FC = Frontal cortex; OC = Occipital cortex.

We explored whether the different PCA groups were associated with clinical phenotypes such as age, dementia status, or MAPT genotype. Group 1 controls without pathology were younger compared to the other groups (Figure 6c) and tau pathology was associated with age in controls (Supplementary Table 1, Supplementary Figure 5). Most of the PD cases with dementia were in Group 4 (Figure 6d), and there was no clear association between MAPT genotype and PCA group (Figure 6e).

## Discussion

In this study, we used single-molecule imaging to compare the distribution of nanoscopic tau aggregates with IHC-labelled tau inclusions in PD post-mortem brain. We identified a unique population of high-intensity nanoscopic tau aggregates in the PD putamen that were not detectable using IHC, ranging from 10–1,000 epitopes per aggregate and up to 1,000 nm in length. Our results align with recently-published results of nigrostriatal tau deposition associated with motor symptom development, as well as implicating soluble tau aggregates in PD-dementia (18).

Although we observed consistently higher αSyn IHC in PD, αSyn SiMPull did not identify a PD-specific nanoscopic αSyn aggregate population. Instead, we observed low numbers of low-intensity αSyn aggregates, predominantly composed of fewer than five epitopes. The lack of association between αSyn SiMPull and IHC forced us to question whether these were true αSyn aggregates, functional low-order assemblies such as tetramers, or alternatively, αSyn monomers sandwiched by other protein binding partners (27, 28). If the latter is true, this may explain the increased αSyn SiMPull intensities in the frontal cortex and occipital cortex compared to other brain regions, as the cortical regions were largely free of Lewy Bodies and may contain more functional αSyn engaged in protein-protein interactions. At this stage we were unable to determine which assemblies were true nanoscopic αSyn aggregates, and future studies should consider screening αSyn binding partners as exclusion criteria for aggregate analysis.

Unlike αSyn, we identified a robust nanoscopic tau aggregate population that aligned with IHC staining in the hippocampus and amygdala, which are highly and early affected in Alzherimer’s disease pathology (13). Surprisingly, we observed a striking increase in SiMPull intensity in the PD putamen in the absence of tau IHC staining. From SiMPull, the number of epitopes per aggregate had a bimodal distribution, with one population of aggregates between 2–10 epitopes and a second between 10–1,000 epitopes. The first tau aggregate population had a similar number of epitopes as αSyn SiMPull, and could similarly be interpreted as true tau aggregates, or tau complexes with binding partners. Further analysis of the underlying intensity distributions for each brain region revealed that Path-Negative controls only had low-intensity (or 2–10 epitope) aggregates, reinforcing that protein assemblies in this size range were not associated with IHC, whilst high-intensity aggregates were observed in the hippocampus and amygdala for Path-Positive controls and PD. In the putamen, high-intensity aggregates were uniquely present in PD, and the substantia nigra displayed a similar trend towards higher intensities in a reduced sample size. The putamen is perturbed in PD due to its functional and anatomical proximity to the substantia nigra, although the trigger for cell death in this circuit is not known (29, 30). The recent study by Chu et al., (2024) proposed that dopaminergic neurodegeneration may be tau-mediated, after observing IHC-positive nigrostriatal tau pathology as a common feature of PD and non-PD individuals with motor deficits. The average age of non-PD motor-deficit individuals in this study was 10 years older than our PD cohort, implying that the nanoscopic tau aggregates observed by SiMPull may eventually form large tau inclusions. Furthermore, our results support the conclusion that neuronal cell death in PD may be tau-mediated, but raise the question of which aggregate species are more neurotoxic.

Our findings in the putamen were strengthened by the results of phosphorylated tau (pTau) SiMPull, which revealed a consistent increase in pTau aggregate intensity in PD. This time, there was a single aggregate intensity distribution for each group and lower numbers of aggregates in Path-Negative controls compared to the other groups, indicating that the low-intensity aggregate population from tau SiMPull was not phosphorylated. Although the occipital cortex had significantly higher pTau aggregate levels in PD, Path-Positive controls and PD had similar intensity distributions, therefore occipital cortex tau pathology did not appear PD-specific. Again, the putamen had significantly higher intensity aggregates in PD compared to both control groups, and we used super-resolution imaging to determine that their lengths ranged from 30–1,000 nm. The differences in putamen pTau aggregate sizes were influenced by a subset of PD cases with mean length greater than 120 nm, and the longest aggregates appeared fibrillar in shape from reconstructed super-resolution images. One limitation of this study is that we were unable to use Thioflavin T with SiMPull to determine if nanoscopic aggregates were beta-sheet positive due to high background fluorescence, and greater structural analysis of nanoscopic tau aggregates would be of interest in the future.

Tau SiMPull and tau IHC aggregates were closely linked in the control group, however, that relationship was disrupted in PD. In PD, tau SiMPull and IHC were only correlated in the hippocampus, but there was no relationship in the amygdala or putamen, despite high levels of bright nanoscopic tau aggregates. Comparing tau SiMPull with αSyn IHC coverage in PD may provide insights into regional differences; there was a positive correlation between tau SiMPull and αSyn IHC in the hippocampus, indicating a universal response by aggregation-prone proteins in this region. In the amygdala, tau SiMPull was negatively correlated with αSyn IHC, indicating a possible interaction between αSyn and tau; for example tau may be incorporated into Lewy Bodies, resulting in the absence of nanoscopic tau aggregates (31). An inherent limitation of working with post-mortem brain tissue is the inability to determine the sequence of pathological events, so it is not possible to determine from this data whether Lewy Bodies affect tau aggregate levels or vice versa. Nevertheless, these results indicate a distinct distribution pattern for tau aggregates in PD compared to controls.

Combining aggregate measurements from SiMPull and IHC prompted two different interpretations for the role of tau pathology in PD. We used principal component analysis to divide the cohort into four clusters: Group 1 controls had low pathology in all brain regions; Group 2 controls had high tau pathology that was associated with advanced age; Group 3 PD had αSyn pathology and moderate tau pathology in the hippocampus and amygdala; Group 4 PD had αSyn pathology and high tau pathology, with a unique high-intensity nanoscopic tau aggregate population in the putamen, and the highest rates of dementia. It should be noted that PD-dementia was not a selection criterion for this cohort, so a follow-up study will determine the strength of this relationship in a larger cohort of PD versus PD-dementia. The different patterns of tau pathology in Groups 3 and 4 can be interpreted as: (1) Tau pathology is independent of PD pathogenesis, beginning in the hippocampus and progressing along the tau Braak stages, therefore Group 4 is more advanced in the process compared to Group 3; (2) Tau aggregation initiates in the basal ganglia in a subset of PD cases like Group 4, and progresses in parallel to age-related tau aggregation that begins in the hippocampus. In the latter scenario, tau aggregates could have a central role in causing dopaminergic cell loss in PD, as previously discussed. Alternatively, tau aggregation may be triggered by cell death in the substantia nigra, for example by reduced dopamination of tau which was recently shown to protect against aggregation (32). If tau is truly driving cell death in PD, it could be important to stratify PD patients based on nigrostriatal tau pathology to identify a subpopulation who would benefit from tau-targeting therapies in the future.

In conclusion, we leveraged single-molecule imaging to measure nanoscopic tau aggregates in PD that may have been previously underestimated by pathological studies relying on IHC. Our data provides evidence that nanoscopic tau aggregate formation in the putamen may contribute to pathology in a subset of cases, and is linked to more advanced PD with dementia. Future studies should aim to establish the sequence of pathological events and the prevalence of nanoscopic putamen tau aggregates in the PD brain, which may represent a targetable aggregate species for disease-modifying therapies.

## Methods

### Case Information

Post-mortem human brain tissue from 14 idiopathic PD cases and 15 control (CON) cases was acquired from the Cambridge Brain Bank. All research procedures utilising human brain tissue were conducted with approval by the London-Bloomsbury Research Ethics Committee (16/LO/0508) and with informed consent from the families of the donors. Idiopathic PD cases received a clinical diagnosis of PD during life and displayed post-mortem Lewy Body neuropathology. Clinical data pertaining to PD progression was available for a subset of PD cases who attended Cambridge University Hospital clinics during life or from death certificates, and Braak staging for αSyn and tau was assessed by neuropathologist Dr Annelies Quaegebeur (Table 1).

Control cases had no diagnosis of PD during life or PD-associated post-mortem neuropathology. However, some controls were documented as having dementia or other neurological disease on their death certifications (Table 1). Additionally, control cases CON5, CON6, CON7, CON8, CON9, CON11, CON12, and CON14 had tau neurofibrillary tangle deposition, although it was limited to the hippocampus and amygdala.

From the available information in the PD group, there was a mean disease duration of (10.28 ± 5.7 years), 5/14 cases had dementia, a range of Lewy Body Braak stages from IV–VI, and a range of tau Braak stages from I–III.

Upon arrival at the Brain Bank, brains were bisected and one half was flash-frozen and stored at −80 °C while the other half was fixed in 10% neutral buffered formalin for 14 days. The formalin-fixed tissue was embedded in paraffin blocks and cut into 10 μm sections for immunohistochemical analysis. Tissue sections from six brain regions were acquired for this study including hippocampus (PD n = 14; control n = 15), amygdala (PD n = 14; control n = 12), substantia nigra (PD n = 13; control n = 15), putamen (PD n = 14; control n = 15), occipital cortex (PD n = 14; control n = 15), and frontal cortex (PD n = 14; control n = 15).

Frozen tissue was acquired for the same cases and brain regions and processed for single-molecule imaging. It should be noted that frozen tissue availability varied for some regions, and the SN in particular was only available for a subset of cases: hippocampus (PD n = 12; control n = 11); amygdala (PD n = 12; control n = 12), substantia nigra (PD n = 5; control n = 4), putamen (PD n = 14; control n = 15), occipital cortex (PD n = 14; control n = 15), and frontal cortex (PD n = 14; control n = 15).

### Immunohistochemistry (IHC)

IHC was performed by the Cambridge Brain Bank as previously described (33). In brief, tissue sections were deparaffined and hydrated by sequential incubation in xylene, ethanol, and water. The sections were pre-treated with formic acid for 5 minutes for additional antigen retrieval. Sections were incubated with hydrogen peroxide for 5 mins, followed by washing with water and phosphate-buffered saline (PBS). Sections were blocked using 2% Marvel in PBS for 20 mins, then incubated with the primary antibody (Table 2) in 2% Marvel for 1 hour. The sections were washed with PBS for 5 mins, then incubated with the secondary antibody (1% Rabbit horse radish peroxidase, diluted in 10% serum in PBS) for 1 hour. The sections were washed with PBS, then incubated with 3,3′-diaminobenzidine (DAB) (Vector, SK-4100). Finally, the sections were counterstained with hematoxylin to visualize cell nuclei, and sealed for imaging.

**Table 2.** Primary antibodies used for IHC.

| Target | Type/Species | Source | Ref | Dilution |
| --- | --- | --- | --- | --- |
| αSyn | Polyclonal/Rabbit | Enzo Life Sciences | BML-SA3400-0100 | 1:250 |
| Tau | Monoclonal/Mouse | Thermo Fisher Scientific | MN1020 | 1:500 |

**Table 3.** Monoclonal antibodies used for SiMPull. Epitope refers to the amino acid sequence of the target protein. AF= Alexa fluor; N/A = Not available.

| Target | ID | Species | Epitope | Source | Ref |
| --- | --- | --- | --- | --- | --- |
| αSyn | Syn211 | Mouse | 121-125 | Santa Cruz | SC-12767 |
| αSyn | Syn211-AF647 | Mouse | 121-125 | Santa Cruz | SC-12767 AF647 |
| αSyn | SOY1 | Mouse | C-terminus | Merck | MABN1818 |
| Tau | Biotin HT7 | Mouse | 159-163 | Invitrogen | MN1000B |
| Tau | HT7 | Mouse | 159-163 | Invitrogen | MN1000 |
| pTau | Biotin AT8 | Mouse | pSer202,<br>Thr205 | Invitrogen | MN1020B |
| pTau | AT8 | Mouse | pSer202,<br>Thr205 | Invitrogen | MN1020 |

The pTau antibody AT8 was also used for super-resolution imaging with DNA points accumulation for imaging in nanoscale topography super-resolution imaging (DNA-PAINT). The antibody was conjugated with the following docking oligonucleotide sequence:
DBCO TEG-AAACCACCACCACCACCACCACCACCACCACCA

### IHC imaging and analysis

Slide scanning at 20 x magnification was performed at the Cambridge Brain Bank or Cancer Research UK Cambridge Institute using the Aperio Scanscope AT2 (Leica Biosystems). Image pre-processing and quantitative analysis was performed using QuPath software (version 0.4.3) as described by a previously published protocol (34). Briefly, colour deconvolution was applied to the scanned brightfield whole slide images to separate the stains into different channels. Slides were then manually inspected to remove pigmented cells (in the substantia nigra), DAB artefacts, or folded tissue, and the region of interest (ROI) was outlined using the wand tool to exclude white matter. A thresholder tool in QuPath was applied to detect DAB-positive pixels (Resolution = Full (µm/pixel), Pre-filter = Gaussian, Smoothing sigma = 0.5, Threshold = 1.3). IHC coverage for each marker was calculated by:

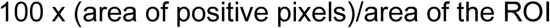

### Frozen post-mortem human brain tissue fractionation

The steps for frozen brain tissue fractionation were adapted from a previously published (20). The case order of brain tissue processing was randomised to avoid technical bias and different dounce homogenisers were used for PD and CON cases to avoid cross-contamination. Briefly, 200-300 mg of fresh-frozen brain tissue was homogenised manually (Sigma-Aldrich, Cat. D8938) in five volumes (w/v) of ice cold low-salt buffer (124 mM NaCl, 2.8 mM KCl, 1.2 5mM NaH_2_PO_4_, 26 mM NaHCO_3_; pH 7.32, supplemented with 5 mM EDTA, 1 mM EGTA, 5 μg/mL leupeptin, 5 μg/mL aprotinin, 2 μg/mL pepstatin, 20 μg/mL Pefabloc, 5 mM NaF), with 10 strokes of a high-pass pestle followed by 15 strokes of a low-pass pestle. The homogenate was centrifuged for 30 min at 21,000 xg at 4 °C and the supernatant was collected. The total protein concentration of the supernatant was measured immediately by Pierce bicinchoninic acid (BCA) protein assay kit (Thermo Scientific, Cat. 23227) according to the manufacturer’s instructions, then aliquoted and stored at −80 °C.

### Antibodies for single molecule pull-down (SiMPull)

Where possible, antibodies were acquired pre-labelled with biotin and Alexa Fluor (AF) dyes. The remaining antibodies were home-labelled as described below.

The complimentary imaging oligonucleotide was conjugated with Cy3b dye, with the sequence: TGGTGGT-AminoC7 – Cy3B

### Antibody labelling

Capture antibodies were biotinylated, and detection antibodies were conjugated with a docking oligonucleotide for DNA-PAINT, using the SiteClick™ Antibody Azido Modification Kit (Invitrogen, Cat. S20026) according to the manufacturer’s instructions. In brief, at least 150 µg of antibody without preservatives or salts was concentrated and buffer exchanged with Buffer A using a 50 kDa MWCO Amicon spin filter (Merck Millipore, Cat. UFC505024). The antibody was recovered and incubated with beta galatosidose overnight at 37 °C. The next day, components E, F, G, and H were combined, and the antibody was added to the mixture. The mixture was incubated overnight at 30 °C. The next day, the mixture was concentrated using a 50 kDa MWCO Amicon spin filter and 10 molar equivalents of biotin were added, followed by another overnight incubation at 37 °C. Finally, excess biotin was removed using a 40 kDa MWCO Zeba Spin Desalting Column (Thermo Scientific, Cat. A57766), the antibody was concentrated using a 50 kDa MWCO Amicon Spin filter, and the concentration was determined by Nanodrop A280.

The Zip Alexa Fluor^TM^ 647 Rapid Antibody Labeling Kit (Invitrogen, Cat. Z11233) was used to label detection antibodies. The antibodies were first concentrated and buffer exchanged with PBS using a 50 kDa MWCO Amicon spin filter. The antibody was recovered and incubated with 1/10 volume 1 M sodium bicarbonate and 10 molar equivalents of the appropriate dye for 1 hour at 37 °C. The reaction was quenched by diluting 1:1 in 1 M Tris pH 8, then excess dye was removed using a 40 kDa MWCO kDa Zeba Spin Desalting Column. The antibody was concentrated using a 50 kDa MWCO Amicon spin filter, then the final concentration and the dye-labelling efficiency were determined by Nanodrop.

### HT7-binding peptide sequence

A dimer of the HT7-binding peptide sequence (KK(PEG)8GAAPPGQKGQ, Cat. No. U9131MEFG0-9, Genscript) was used for tau SiMPull, as previously described (25). The dimer contains two binding sites for the total-tau antibody HT7; when used for SiMPull, the dipeptide may bind the capture antibody and a single detection antibody.

### Silica nanoparticle (SiNaP) aggregate standards

Tau SiNaP aggregate standards were prepared according to the previously published protocol by Böken et al., (2024), by conjugating the HT7 antibody binding sequence to the SiNaPs, and αSyn SiNaP aggregate standards were prepared according to the previously published protocol by Herrmann et al., (2017) (36, 37).

### Coverslip passivation for single-molecule pull-down

Glass coverslips (26 x 76 mm, Epredia) were cleaned using an argon plasma cleaner (PDC-002, Harrick Plasma) for 30 minutes, then a 50-well PDMS gasket (Grace Bio-Labs, Ref. 103250) was attached to each coverslip. The wells were loaded with 6 µL of coating solution (1:1 mixture of Rain-X and isopropanol, filtered using a 0.22 µm PVDF filter (Millex, Ref. SLGV004SL)), and left to evaporate at room temperature as previously described (38). The surfaces were always prepared the day before a SiMPull experiment.

### Single-molecule pull-down (SiMPull)

Tris-buffered saline (TBS, Fisher Bioreagents, Cat. BP2471-500) and TBS-Tween (TBS-T, Rockland antibodies and assays, Cat. MB-013) buffers were filtered using a 0.2 µm PVDF filter (Millex, Ref. SLGP033RS) immediately prior to use. A 0.22 µm PVDF filter (Millex, Ref. SLGV004SL) was used for all other reagents that are described as filtered. All reagents or wash buffers were added to the wells in 10 µL volumes, and all incubations were carried out at room temperature.

Pre-prepared Rain-X coated coverslips were placed in a humidity chamber to prevent the surface from drying out. The wells were washed twice with TBS (Gibco, Ref. 10010-015), then incubated with NeutrAvdin (Thermo Scientific, Ref. 31050) at 0.1 mg/mL in TBS for 15 minutes. The wells were washed three times with TBS, then incubated for 1 hour with 1% pluronic F-127 polymer (Biotium, Cat. 59005, 10% in water) diluted 1:10 in TBS and filtered. The wells were washed three times with TBS-T (1x PBS with 0.05% Tween 20; Rockland, Code MB-075-1000), then incubated with 10 nM biotinylated Capture antibody for 15 minutes. The wells were washed twice with TBS-T, then incubated with filtered blocking solution (500 µL TBS-T, 50 µL BSA (R&D Systems, Cat. 841380), 20 µL normal goat serum (Abcam, Cat. Ab7481), 20 µL recombinant BSA (New England BioLabs, Cat. B9200S)), for 30 minutes to prevent non-specific protein adsorption to the surface. The wells were subsequently washed twice with TBS-T before sample addition.

For tau and pTau SiMPull: brain samples were diluted 2-fold in TBS and incubated for 2 hours. For αSyn SiMPull: brain samples were added to the surface without dilution and incubated for 3 hours. Samples were run in triplicate for each brain region and antibody pair, with one well per case used on each surface. For every surface, one well was incubated with TBS (negative control), and another was incubated with a single-detection antibody control (HT7-dipeptide for tau SiMPull, capture and detect with HT7; αSyn monomer for αSyn SiMPull, capture with SOY1 and detect with Syn211). Following sample incubation, the wells were washed four times with TBS-T.

For diffraction-limited imaging, the wells were incubated with 5 nM detection antibody labelled with Alexa-Fluor 647 in filtered 1% BSA in TBS-T for 15 minutes. For multichannel imaging, both detection antibodies were added at a final concentration of 5 nM. Finally, the wells were washed three times with TBS-T, then the buffer was exchanged with TBS for image acquisition.

For super-resolution imaging, another blocking step was performed with filtered 1% BSA in TBS-T for 30 minutes. Next, the DNA-conjugated detection antibody diluted in TBS was incubated for 15 minutes. The wells were washed twice with TBS-T, then 1 nM Cy3b-conjugated imaging strand was added to the wells, and a coverslip was added on top to prevent evaporation throughout imaging.

### Diffraction-limited image acquisition and analysis

Diffraction-limited imaging was performed using a home-built total internal reflection fluorescence (TIRF) microscope using an inverted Ti-E Eclipse body (Nikon Corporation) fitted with a 1.49 N.A, 100x objective (Apo TIRF, Nikon) and in-built Perfect Focus System. The microscope was fitted with 488 nm (iBeam – SMART, Toptica) and 638 nm lasers (MLD, 638, Cobalt), and the fluorescent emissions was collected by an EMCCD camera (Evolve 512, Photometrics) through a 1.5x beam expander, with an electron-multiplying gain of 214 ADU/photon, sensitivity of 3.019 electrons/ADU, and offset of 604.2 ADU. Each pixel corresponded to 109.2 nm and image acquisition (50 frames, 50 ms exposure time, 12 fields of view) was controlled with MicroManager (39).

Automated analysis of single-molecule localisation was performed in ImageJ (40). The code was adapted from https://github.com/YPZ858/DF-single-molecule-couting-/blob/main/DF21.ijm. Briefly, the image frames 10–50 were combined into a z-stack, then the ThunderSTORM plugin was used to apply a wavelet filter of 2*std(Wave.F1) and detect single molecules (41). The results were exported as an Excel spreadsheet and further filtering was performed to retain data that met the following criteria: higher intensity than the mean intensity of the blank (TBS only); replicates with positive signal in at least 9 out of 12 fields of view; cases with positive signal in at least 2 out of 3 replicates.

Example images were prepared in ImageJ by first performing a z-stack of frames 10–50. The greyscale values were calibrated to photons using the ThunderSTORM plugin, and images were contrast-adjusted with a minimum value of 0 and a maximum value corresponding to the range of spot intensities in the image. The calibration bar was used to demonstrate the contrast ranges of different images, and a higher maximum value corresponded to higher intensity spots. It should be noted that the maximum value of the calibration bar does not necessarily indicate the maximum photon value in the image, as contrast ranges were selected as a compromise between multiple images being compared. The field of view was cropped to half of the original size to enlarge the image sufficiently for demonstration purposes.

### Intensity calibration to calculate number of epitopes per aggregate

The intensity of a single detection antibody was used to calculate the approximate number of epitopes per aggregate. To obtain this intensity value, HT7 dipeptide and αSyn monomers (Abcam, cat. AB218818) were used for tau and αSyn SiMPull respectively (Methods Figure 1). To account for the epitope bound by the capture antibody, +1 was added.

### Super-resolution image acquisition and analysis

Super-resolution imaging was performed with DNA-PAINT using the same optical setup as diffraction-limited imaging. Images were acquired using the 561 nm laser (Jive, Cobalt), with 8,000 frames, 100 ms exposure time, and 4 fields of view.

DNA-PAINT image reconstruction and analysis was performed using the Picasso package, as previously described (25, 42). The custom scripts are available at https://zenodo.org/records/8027256. In brief, the first 300 frames were discarded, localisations were identified and fitted, using redundant cross-correlation to correct for microscope drift. The localisations were filtered using a 30 nm precision threshold, then assigned to clusters with DBSCAN using the scitkit-learn package (radius of 2 and minimum density of 1, minimum of 3 localisations per cluster) corresponding to individual aggregates (43). Each aggregate was measured in terms of skeletonised length using SKAN, calculated as the summed branch distance (44). The results were exported as an Excel spreadsheet, and were further filtered to retain clusters that had a higher number of localisations compared to the blank (TBS only).

### DNA extraction and MAPT genotyping

DNA was isolated from fresh-frozen brain tissue using the QIAamp DNAMini and Blood Mini kit according to the manufacturer’s instructions. H1 and H2 MAPT haplotype was determined by Taqman assay (Assay ID: C 7563692_10; SnP ID: rs1800547) using the QuantStudio 5 detection system (Applied Biosystems).

### Statistical analysis in R

All statistical analysis was performed in R (Version 4.2.2) and all graphs were plotted using the packages ggplot2, tidyverse, plyr, ggpubr, scales, ggh4x, ggrepel (45–52). The appropriate statistical test was determined after checking the data met the normality assumptions, and p-values of less than 0.05 were considered significant. The sample size for all whole-cohort comparisons was PD n = 14 and CON n = 15. For comparisons of IHC coverage, mean SiMPull density or mean SiMPull intensity between PD and CON, non-parametric two-sided Mann-Whitney test was used. For linear correlation analysis of human samples, data was not normally distributed therefore the non-parametric Spearman correlation was performed using stat_cor in the R package ggpubr, and the correlation coefficient rho (ρ) was displayed. For linear correlation analysis of SiNaP SiMPull, data was normally distributed, therefore the Pearson correlation was performed using stat_cor in the R package ggpubr, and the correlation coefficient R^2^ was displayed. Box plots show the first quartile to third quartile (box) with the median marked, and whiskers extend to 1.5 times the interquartile range. For intensity distribution histograms, intensity was plotted on a log_10_ x-axis scale, and the y-axis was plotted as percentage per group to account for the different sample sizes. For multiple comparisons of age of death, non-parametric Dunn’s multiple comparison test with Benjamini-Hochberg p-value adjustment was used.

The following input variables were used for principal component analysis (PCA): αSyn IHC coverage (%) for hippocampus, amygdala, putamen, substantia nigra; Tau IHC coverage (%) for hippocampus, amygdala, putamen, substantia nigra; Tau SiMPull mean bright aggregates (%) for hippocampus, amygdala, putamen; pTau SiMPull mean intensity (photons) for hippocampus, amygdala, putamen. Results from the frontal cortex and occipital cortex were excluded due to low signal for all markers, and SiMPull results from the substantia nigra were excluded due to low sample size. For each variable, missing were imputed with the median, and data was centred and scaled prior to PCA. Subsequently, k-means cluster analysis was performed using principal components 1 and 2. For comparisons between PCA groups for age of death, dementia status, or MAPT genotype: Group 1 n = 7; Group 2 n =8; Group 3 n = 9; Group 4 n = 5.

## Acknowledgements

We gratefully acknowledge the contributions of our brain tissue donors and their families and the Cambridge Brain and Tissue Bank. We acknowledge the histology core at the Cancer Research UK Cambridge institute for their technical support. The work was supported by a grant from the UK Medical Research Council.

## Author Contributions

F.L.: Tissue processing, single-molecule assays, immunohistochemistry image analysis, data analysis, manuscript preparation, conception and design. D.B.: Development of tau assays, super-resolution data analysis. Y.Z.: Assay development, technical support. K.H.: Tissue preparation, immunohistochemistry. D.R.: Tissue preparation, immunohistochemistry. L.K.: Tissue preparation, technical support. B.P.: Data collection. A.Q.: Neuropathological assessment, manuscript preparation. C.H.W.G.: Manuscript preparation, conception and design. D.K.: Manuscript preparation, conception and design, supervision. The final manuscript was edited and approved by all authors.

