## Supplementary for "Nanoscopic tau aggregates in Parkinson’s disease"

### Nanoscopic tau aggregates in Parkinson's disease: Coincidence or co-pathology?

#### Methods

##### Western blot

Brain samples were diluted to 15 µL containing 25 µg total protein in 4x Laemmli protein sample loading buffer (Bio-rad, Cat. 1610747) containing 1:10 2-mercaptoethanol (Bio-rad, Cat. 1610710). The samples were boiled at 70 °C for 10 minutes and ran at 100 V on a 4-20% Mini-PROTEAN TGX precast gel (Bio-rad, Cat. 4561096) or 4–15% gradient Mini-PROTEAN TGX precast gel (Bio-rad, Cat. 4568086) for detecting αSyn and tau respectively. The gel was transferred onto a 0.2 µm nitrocellulose membrane (Bio-rad, Cat. 1620146) using the TurboBlot (Bio-rad), the membrane was fixed using 4% paraformaldehyde, blocked with 5% non-fat milk, then incubated with the primary antibody Syn211 or HT7 overnight at 4 °C. The next day, the membrane was incubated with the HRP-conjugated anti-mouse secondary antibody (Promega, Cat. W4021) for 1 hour at room temperature, followed by the ECL substrate (Bio-rad, Cat. 170-5061). The membrane was visualised using the ChemiDoc MP Imaging System (Bio-rad). The membrane was stripped and re-probed with GAPDH (Thermo Scientific, Cat. MA515738) as a loading control.

**Antibodies for single-molecule pull-down**

**Supplementary Methods Table 1. Monoclonal αSyn antibodies used for SiMPull.** Epitope refers to the amino acid sequence of the target protein. AF647 = Alexa fluor 647.

| Target | ID | Species | Epitope | Source | Ref |
|---|---|---|---|---|---|
| αSyn | 5G4 | Mouse | 44-57 | Merck | MABN389 |
| αSyn | Biotin LB509 | Mouse | 115-122 | BioLegend | 807710 |
| αSyn | LB509 | Mouse | 115-122 | BioLegend | 807701 |
| αSyn | Biotin MJFR14 | Rabbit | Conformation-specific | Abcam | ab227047 |
| αSyn | AF647 MJFR14 | Rabbit | Conformation-specific | Abcam | ab216309 |
| αSyn | Syn1 | Mouse | 15-123 | BD Biosciences | 610786 |

#### Results

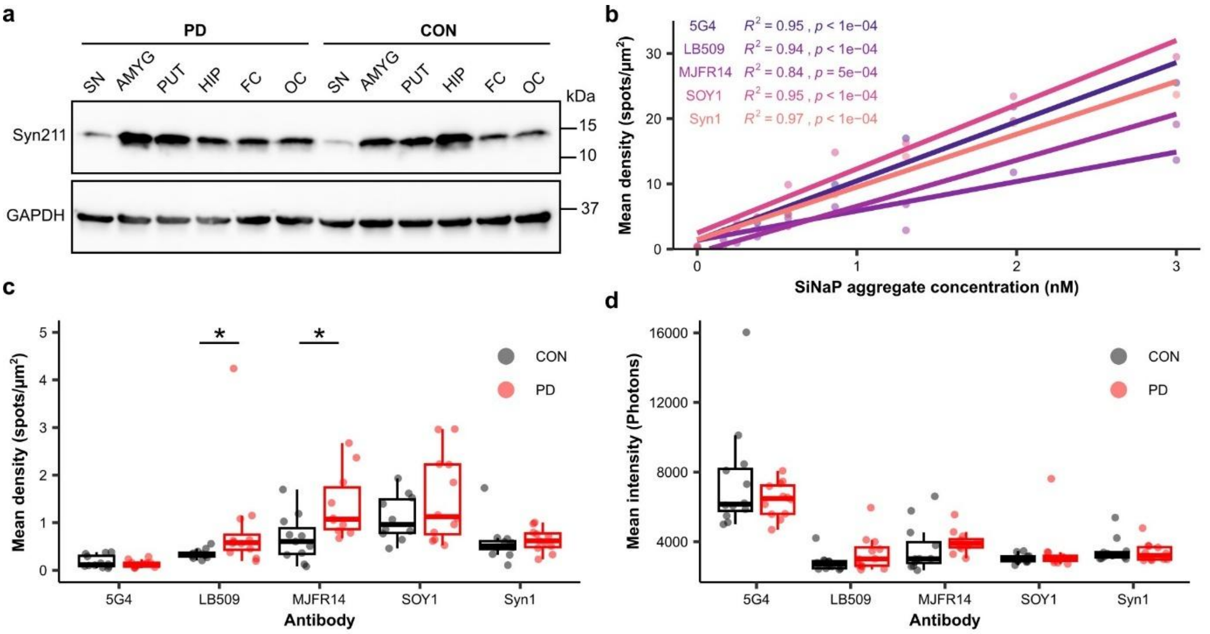


**Supplementary Figure 1. Syn211 binds αSyn in brain samples and similar SiMPull results are obtained using alternative αSyn antibodies. (a)** Western blot using αSyn antibody Syn211 and fractionated brain samples from one PD and one CON case, with a single band visible at 14 kDa corresponding to the molecular weight of αSyn. **(b)** Dilution series of αSyn SiNaP aggregates using five alternative αSyn antibodies. There was a significant relationship between SiNaP concentration and aggregate density for all five antibodies (Pearson correlation). **(c–d)** αSyn SiMPull using AMYG brain samples and five alternative αSyn antibodies. There was a significant increase in **(c)** aggregate density in PD when LB509 and MJFR14 were used (Two-sided Mann-Whitney test, 5G4 $p$ = 0.6297; LB509 $p$ = 0.01004; MJFR14 $p$ = 0.01209; SOY1 $p$ = 0.9774; Syn1 $p$ = 0.1748). **(d)** There were no significant differences in intensity (Two-sided Mann-Whitney test, 5G4 $p$ = 0.7553; LB509 $p$ = 0.16; MJFR $p$ = 0.05966; SOY1 $p$ = 0.7987; Syn1 $p$ = 0.5137). CON = Control; FOV = Field of view; HIP = Hippocampus; AMYG = Amygdala; PUT = Putamen; SN = Substantia nigra; FC = Frontal cortex; OC = Occipital cortex.

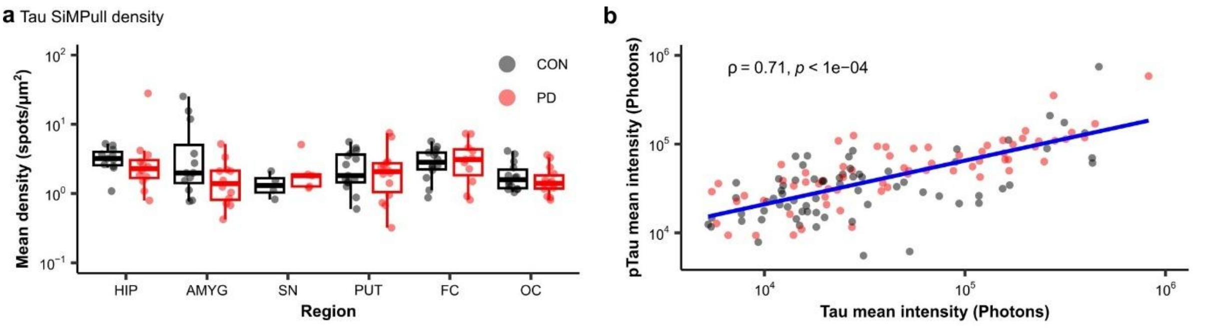


**Supplementary Figure 2. Tau SiMPull densities and relationship between Tau and pTau SiMPull intensities. (a)** There were no significant differences in aggregate density for tau SiMPull between PD and CON (Two-sided Mann-Whitney, HIP $p$ = 0.1037; AMYG $p$ = 0.1978; SN $p$ = 0.4127; PUT $p$ = 0.683; FC $p$ = 0.8931; OC $p$ = 0.5714). **(b)** Tau SiMPull and pTau SiMPull mean aggregate intensities were positively correlated (Spearman correlation). **(a–b)** Each point represents the mean of 9–12 FOVs and 2–3 independent repeats per case. CON = Control; HIP = Hippocampus; AMYG = Amygdala; PUT = Putamen; SN = Substantia nigra; FC = Frontal cortex; OC = Occipital cortex.

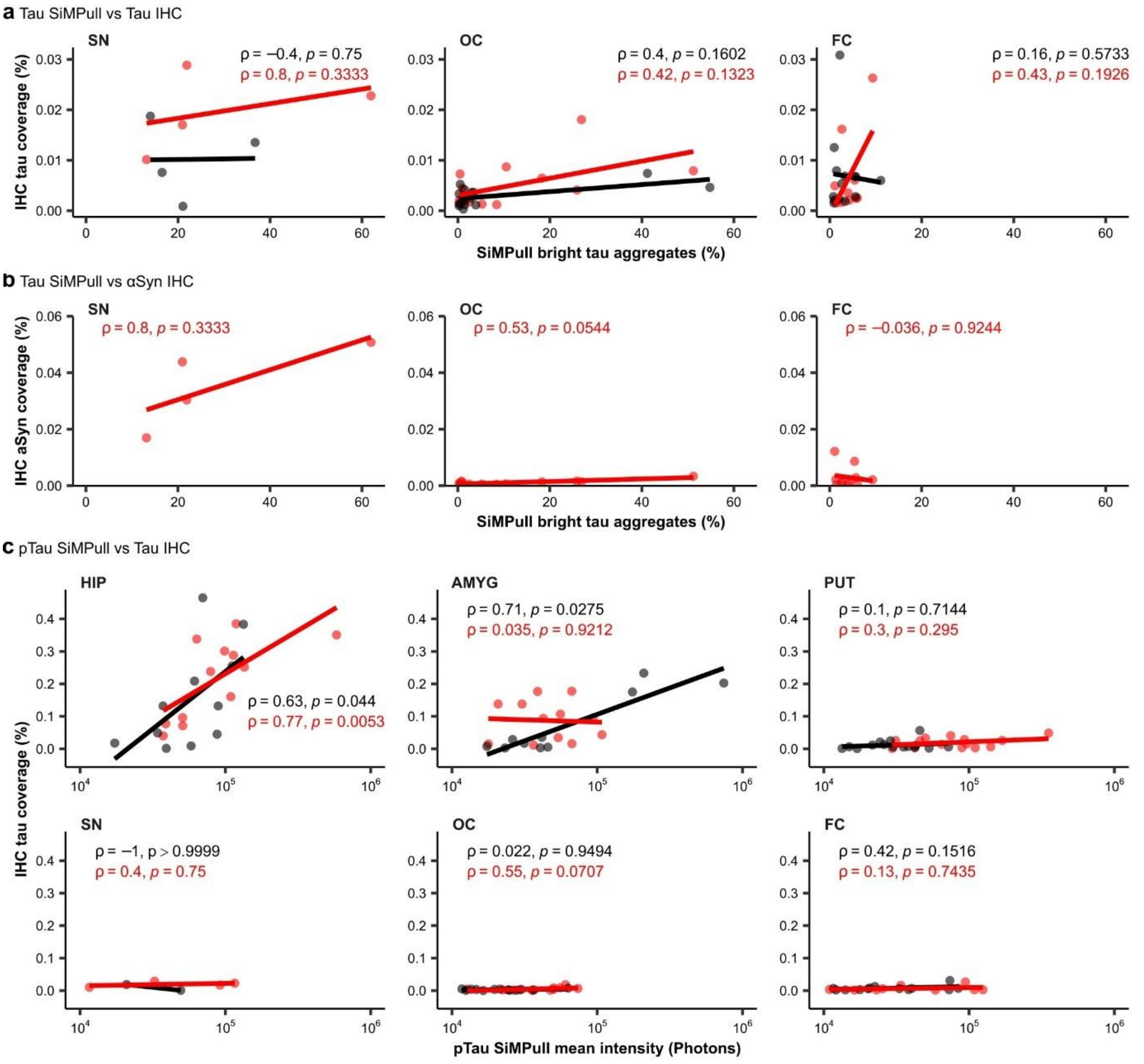

**Supplementary Figure 3. (a)** Tau SiMPull bright aggregates (%) were not correlated with tau IHC in the SN, FC or OC (Spearman correlation). **(b)** Tau SiMPull bright aggregates were not correlated with αSyn IHC in the SN, FC, or OC (Spearman correlation). **(c)** pTau SiMPull mean aggregate intensities were correlated with tau IHC in CON HIP and AMYG, as well as PD HIP (Spearman correlation). **(a–c)** For SiMPull, each point represents the mean of 9–12 FOVs and 2–3 independent repeats per case. HIP = Hippocampus; AMYG = Amygdala; PUT = Putamen; SN = Substantia nigra; FC = Frontal cortex; OC = Occipital cortex.

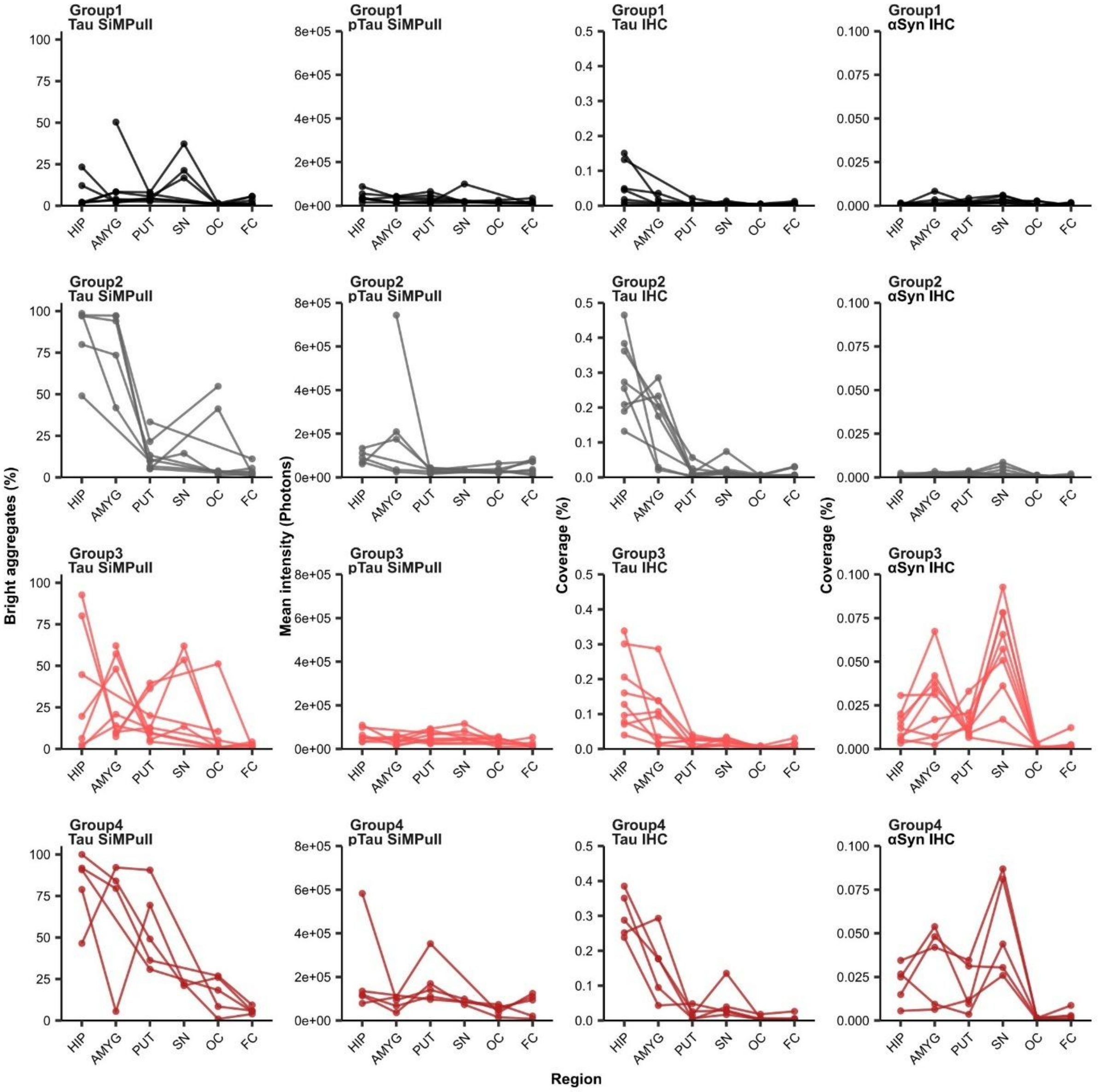


**Supplementary Figure 4. Soluble and insoluble pathology trends in PCA groups.** For SiMPull, each point represents the mean of 9–12 FOVs and 2–3 independent repeats per case. For IHC, each point represents the coverage per case. CON = Control; HIP = Hippocampus; AMYG = Amygdala; PUT = Putamen; SN = Substantia nigra; FC = Frontal cortex; OC = Occipital cortex.

**Supplementary Table 1. Spearman correlation between pathology markers and age of death.** Tau SiMPull was expressed as the percentage of bright tau aggregates, αSyn and pTau SiMPull was expressed as mean intensity, tau IHC and αSyn IHC were expressed as area coverage positive for stain. Control and PD cases were pooled for correlation analysis. Statistically significant correlations were highlighted green. HIP = Hippocampus; AMYG = Amygdala; PUT = Putamen; SN = Substantia nigra; FC = Frontal cortex; OC = Occipital cortex.

| Age of death | | | |
|---|---|---|---|
| **Marker** | **Region** | **ρ** | **p-value** |
| Tau SiMPull | HIP | 0.3575 | 0.0940 |
| | **AMYG** | **0.6659** | **0.0004** |
| | SN | 0.1092 | 0.7796 |
| | PUT | 0.3302 | 0.0802 |
| | FC | 0.2120 | 0.3089 |
| | **OC** | **0.5424** | **0.0029** |
| pTau SiMPull | HIP | 0.2174 | 0.3191 |
| | **AMYG** | **0.4777** | **0.0183** |
| | SN | 0.3698 | 0.3274 |
| | PUT | 0.0624 | 0.7476 |
| | **FC** | **0.4711** | **0.0175** |
| | OC | 0.3392 | 0.0901 |
| Tau IHC | HIP | 0.2833 | 0.1364 |
| | **AMYG** | **0.5450** | **0.0040** |
| | SN | 0.3252 | 0.0912 |
| | **PUT** | **0.3684** | **0.04921** |
| | FC | 0.2937 | 0.1220 |
| | OC | 0.0896 | 0.6440 |
| αSyn SiMPull | HIP | 0.2679 | 0.2165 |
| | AMYG | 0.0889 | 0.6795 |
| | SN | -0.4202 | 0.2602 |
| | PUT | -0.0415 | 0.8309 |
| | FC | 0.0168 | 0.9311 |
| | OC | 0.0244 | 0.8999 |
| αSyn IHC | HIP | 0.1518 | 0.4319 |
| | AMYG | 0.0295 | 0.8864 |
| | SN | 0.1382 | 0.4830 |
| | PUT | 0.3467 | 0.0654 |
| | FC | -0.1463 | 0.4487 |
| | OC | 0.2808 | 0.1400 |

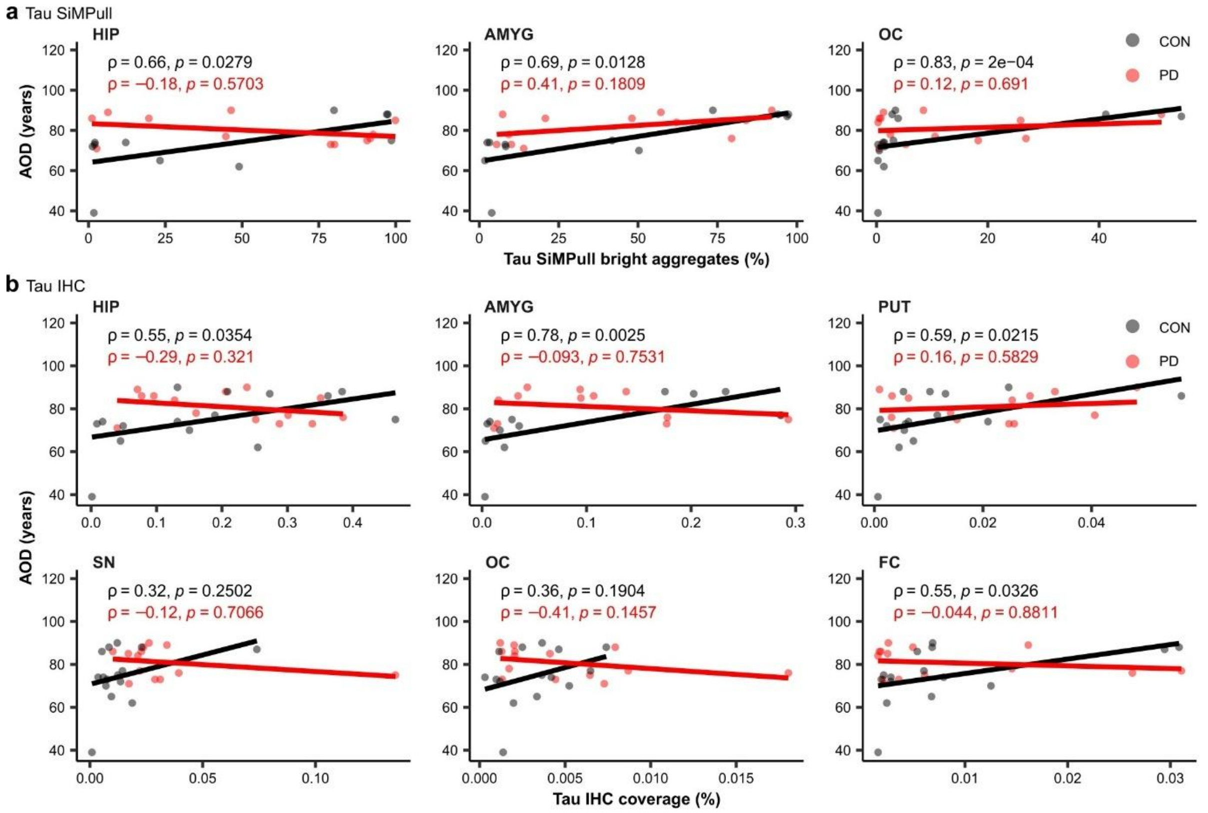


**Supplementary Figure 5. Age of death (AOD) correlates with tau SiMPull and tau IHC in a subset of regions. (a)** Tau SiMPull was correlated with AOD only in CON in the HIP, AMYG, and OC (Spearman correlation). **(b)** Tau IHC was positively correlated with AOD in CON HIP, AMYG, PUT, and FC (Spearman correlation). Tau IHC was correlated with AOD in PD OC only (Spearman correlation). Each point represents the mean per case. CON = Control; HIP = Hippocampus; AMYG = Amygdala; PUT = Putamen; SN = Substantia nigra; FC = Frontal cortex; OC = Occipital cortex.
